## Supplementary material for "Topological Analysis of Differential Effects of Ketamine and Propofol Anesthesia on Brain Dynamics": scripts and csv files: analysis.py

```
#!/usr/bin/env python3
# -*- coding: utf-8 -*-
"""
Created on Tue Jun 18 19:06:49 2019

@author: thosvarley
"""
import numpy as np 
import pandas as pd
import os 
from pathlib import Path 
import matplotlib.pyplot as plt
from ripser import ripser
from persim import plot_diagrams
from collections import Counter
from scipy.stats import entropy
from nolds import lyap_e
from scipy.spatial.distance import pdist, squareform
from scipy.signal import decimate
import sys
sys.path.append("/home/thosvarley/Code/OPyN")
from opyn import optimal_lag, OPN

sys.path.append("/home/thosvarley/Desktop/opn_emergence/scripts/causal_emergence/causal_clustering")
from causal_clustering import determinism_mat, degeneracy_mat, determinism_graph, degeneracy_graph, spectral_clustering, macro

cwd = '/home/thosvarley/Desktop/cuts_stationary/scripts'
ptd = str(Path(cwd).parent)
epc_dir = ptd + "/processed/filtered/"
opn_dir = ptd + "/processed/ordinal_partition_network/"
out_dir = ptd + "/results/"

optimal_dim = 5

PHom_anal = False
OPN_anal = True

plot = True

PH_measures = {}
OPN_measures = {}

def number_cycles(dgm1,thr):
    return ((dgm1<=thr).sum(axis=1)==1).sum()
            
def normalized_persistence_entropy(diagram):
    from collections import Counter
    from scipy.stats import entropy
    return entropy(list(Counter(diagram[:,1]-diagram[:,0]).values()),base=2)/np.log2((diagram[:,1]-diagram[:,0]).sum())

def periodicity_score(diagram,G):
    '''
    Parameters
    ==========
    diagram: persistent diagram
    gg: ordinal partition network

    Results
    =======
    float: periodicity score

    '''
    Ln = len(G.vs())/3 -1
    return 1-(np.diff(diagram[1]).T.max()/Ln)

def permutation_entropy(series):
    l = len(series)
    dim = len(series[0].split())
    C = Counter(series)
    
    for c in C:
        C[c] /= l - dim + 1
    
    return entropy(list(C.values()))
    
for file in os.listdir(epc_dir):
    if file.endswith("filtered.npy") and "Propo" in file:
        name = file[:-13]
        
        epc = np.load(epc_dir + file)
        filt = np.load(epc_dir + file)
        
        ###
        # Persistent Homology on Embedded Point Cloud
        ###
        
        if PHom_anal == True:
            
            print("Phom: {0}".format(name))
            
            epc = decimate(epc, 2)
            dmat = squareform(pdist(epc.T, metric="cosine"))
            
            r = ripser(dmat, coeff=2, do_cocycles=False, distance_matrix=True)
            diagrams = r['dgms']
            np.save(out_dir + "diags/phom_diags/" + name +'_phom_diags.npy', diagrams)
            
            if plot == True:
                
                plt.subplots()
                dgm1 = diagrams[1]
                idx = np.argmax(dgm1[:, 1] - dgm1[:, 0])
                plot_diagrams(diagrams, show = False)
                plt.scatter(dgm1[idx, 0], dgm1[idx, 1], 20, 'k', 'x')
                plt.title("Max 1D Birth = {0}, Death = {1} \n {2}".format(round(dgm1[idx, 0],2),
                          round(dgm1[idx, 1],2),
                          name))
                plt.savefig(out_dir + "diags/phom_diags/phom_{0}.pdf".format(name), dpi=150)
                plt.close()
            
            d = diagrams
            PH_measures[name] = {}
            
            PH_measures[name]['max_pers'] = np.max(d[1][:,1]-d[1][:,0])
            PH_measures[name]['ncycle'] =  d[1].shape[0]

            A = d[1].flatten()
            A.sort()
            betti_curve = np.array([number_cycles(d[1],thr) for thr in A])

            PH_measures[name]['max_ncycles'] = betti_curve.max()
            PH_measures[name]['max_ncycles_thr'] = A[betti_curve.argmax()]
            PH_measures[name]['npe'] = normalized_persistence_entropy(d[1])
            PH_measures[name]['condition'] = name[6:11]
            
            if name[:5] in {"PF731", "PF803", "KT810", "KT724"}:
                PH_measures[name]["monkey"] = "George"
            else: 
                PH_measures[name]["monkey"] = "Chibi"
        ###
        # Making Ordinal Partition Networks
        ### 
        
        if OPN_anal == True:
            import igraph as ig
            
            for R in range(filt.shape[0]):
                
                row = filt[R]
                
                lag = optimal_lag(row, 4500)
                if lag < row.shape[0] / (optimal_dim-1):
                    G, series = OPN(row, dim=optimal_dim, lag=lag)
                else:
                    continue
                
                if G.vcount() > 1:
                        
                    series = np.array(series)
                    adjmat = np.array(list(G.get_adjacency(attribute="weight")), dtype="int")
                    
                    np.save(opn_dir + name + "_r" + str(R) + '_opn_adjmat.npy', adjmat)
                    np.save(opn_dir + name + '_opn_series.npy', np.array(series))
                    
                    namerow = name + "_r" + str(R)
                    
                    OPN_measures[namerow] = {}
                    OPN_measures[namerow]['name'] = name
                    OPN_measures[namerow]['condition'] = name[6:11]
                    OPN_measures[namerow]['row'] = R 
                    OPN_measures[namerow]['opt_lag'] = lag
                    OPN_measures[namerow]['nodes'] = G.vcount()
                    OPN_measures[namerow]['edges'] = G.ecount()
                    OPN_measures[namerow]['in-degree'] = np.mean(G.degree(mode="in"))
                    OPN_measures[namerow]['out-degree'] = np.mean(G.degree(mode="out"))
                    OPN_measures[namerow]['var_in-degree'] = np.var(G.degree(mode="in"))
                    OPN_measures[namerow]['var_out-degree'] = np.var(G.degree(mode="out"))
                    
                    adjmat = np.array(list(G.get_adjacency(attribute="weight")))
                    for out in range(adjmat.shape[0]):
                        total = np.sum(adjmat[out])
                        if total != 0:
                            adjmat[out] /= total
                    OPN_measures[namerow]["determinism"] = determinism_mat(adjmat, normed=True)
                    OPN_measures[namerow]["degeneracy"] = degeneracy_mat(adjmat, normed=True)
                    OPN_measures[namerow]["effectiveness"] = OPN_measures[namerow]["determinism"] - OPN_measures[namerow]["degeneracy"]
                    comm = G.community_infomap(edge_weights="weight")
                    OPN_measures[namerow]["modularity"] = comm.q
                    
                    C = Counter(series)
                    S = Counter(C.values())
                    OPN_measures[namerow]['max_frequency'] = max(C.values())
                    OPN_measures[namerow]['var_frequency'] = np.var(list(C.values()))
                    OPN_measures[namerow]['single_visits'] = S[1]/sum(S.values())
                    
                    size = G.vcount()
                    
                    dist = np.array(G.shortest_paths())
                    r = ripser(dist, distance_matrix=True)
                    diagrams = r['dgms']
                    d = diagrams
                    np.save(out_dir + "diags/opn_diags/" + namerow +'_opn_phom_diags.npy', d)
                
                    if plot == True and len(diagrams[1]) > 0:
                        
                        plt.subplots()
                        dgm1 = diagrams[1]
                        idx = np.argmax(dgm1[:, 1] - dgm1[:, 0])
                        plot_diagrams(diagrams, show = False)
                        plt.scatter(dgm1[idx, 0], dgm1[idx, 1], 20, 'k', 'x')
                        plt.title("Max 1D Birth = {0}, Death = {1} \n {2}".format(round(dgm1[idx, 0],2),
                                  round(dgm1[idx, 1],2),
                                  namerow))
                        plt.savefig(out_dir + "diags/opn_diags/opn+phom_{0}.pdf".format(namerow))
                        plt.close()
                    
                    if len(d[1][:,1]-d[1][:,0]) == 0:
                        OPN_measures[namerow]['max_pers'] = np.nan
                    else:
                        OPN_measures[namerow]['max_pers'] = np.max(d[1][:,1]-d[1][:,0])
                    
                    OPN_measures[namerow]['ncycle'] =  d[1].shape[0]
                    OPN_measures[namerow]['ncycle_norm'] = d[1].shape[0]/size
                    
                    if len(d[1]) > 0:
                        A = d[1].flatten()
                        A.sort()
                        betti_curve = np.array([number_cycles(d[1],thr) for thr in A])
                        
                        OPN_measures[namerow]['max_ncycles'] = betti_curve.max()
                        OPN_measures[namerow]['max_ncycles_thr'] = A[betti_curve.argmax()]
                        OPN_measures[namerow]['npe'] = normalized_persistence_entropy(d[1])
                        OPN_measures[namerow]['periodicity'] = periodicity_score(d, G)
                    else:
                        OPN_measures[namerow]['max_ncycles'] = np.nan
                        OPN_measures[namerow]['max_ncycles_thr'] = np.nan
                        OPN_measures[namerow]['npe'] = np.nan
                        OPN_measures[namerow]['periodicity'] = np.nan 
                    
                    OPN_measures[namerow]['lyapunov'] = lyap_e(row)[0]
                    OPN_measures[namerow]["perm_ent"] = permutation_entropy(series)
                    
                    if name[:5] in {"PF731", "PF803", "KT810", "KT724"}:
                            OPN_measures[namerow]["monkey"] = "George"
                    else: 
                            OPN_measures[namerow]["monkey"] = "Chibi"
                    
                    print(namerow)
                else:
                    continue
                
        print(name)

if PHom_anal == True:
    PH_df = pd.DataFrame(PH_measures).T
    PH_df.to_csv(out_dir + "PHom_results", sep = ",")

if OPN_anal == True:
    OPN_df = pd.DataFrame(OPN_measures).T
    OPN_df.to_csv(out_dir + "OPN_result_d5_Propo.csv", sep = ",")
```
