## Supplementary material for "Topological Analysis of Differential Effects of Ketamine and Propofol Anesthesia on Brain Dynamics": scripts and csv files: distance_timeseries.py

```
#!/usr/bin/env python3
# -*- coding: utf-8 -*-
"""
Created on Mon Sep 16 10:35:17 2019

@author: thosvarley
"""
import numpy as np 
import pandas as pd 
from scipy.spatial.distance import pdist, squareform 
import os 

path = "/home/thosvarley/Desktop/cuts_stationary/processed/injection/filtered/"
out = "/home/thosvarley/Desktop/cuts_stationary/processed/injection/filtered/cosine_series/"

max_awake, max_ketam, max_propo = [], [], []
max_v_awake, max_v_ketam, max_v_propo = [], [], []

max_dist = (max_awake, max_ketam, max_propo)
max_velocity = (max_v_awake, max_v_ketam, max_v_propo)

for file in os.listdir(path):
    if file.endswith(".npy"):
        name = file[:21]
        series = np.load(path + file).T
        
        if "Awake" in file:
            condition = 0
        elif "Ketam" in file:
            condition = 1
        else:
            condition = 2
        N = series.shape[0]
        
        d_mat = squareform(pdist(series, metric = "cosine"))**-1
        np.fill_diagonal(d_mat, 0)
        
        for i in range(N):
            d_mat[i] /= np.sum(d_mat[i])
        
        prob_series = np.diagonal(d_mat, offset = 1)
        bin_prob_series = np.zeros(N)
        
        for i in range(N-1):
            if prob_series[i] == max(d_mat[i]):
                bin_prob_series[i] = 1
        
        d_series = np.zeros(N-1)
        
        for i in range(N-1):            
            d_series[i] = pdist(series[i:i+2], metric = "cosine")[0]
        
        max_dist[condition].append(max(d_series))
        max_velocity[condition].append(max(np.diff(d_series)))
        
        np.save(out + name + "_cosine_series", d_series)
```
