## Supplementary material for "Topological Analysis of Differential Effects of Ketamine and Propofol Anesthesia on Brain Dynamics": scripts and csv files: figures.py

```
#!/usr/bin/env python3
# -*- coding: utf-8 -*-
"""
Created on Sun Oct  4 11:25:21 2020

@author: thosvarley
"""
import numpy as np 
import pandas as pd 
import matplotlib.pyplot as plt 
import seaborn as sns 
sns.set(style="darkgrid")
palette = sns.husl_palette(3, l=.4)
light_palette = sns.husl_palette(3, l=.75)
in_dir = '/home/thosvarley/Desktop/cuts_stationary/results/'

opn = pd.read_csv(in_dir + "OPN_results.csv", index_col="Unnamed: 0")
phom = pd.read_csv(in_dir + "PHom_results.csv", index_col="Unnamed: 0")

phom_awake = phom[phom["condition"] == "Awake"]
phom_ketam = phom[phom["condition"] == "Ketam"]
phom_propo = phom[phom["condition"] == "Propo"]

# %%
plt.figure(figsize=(12,8))
metrics = ["max_pers", "ncycle", "max_ncycles"]
titles = ["Maximum Persistence", "Number of Cycles", "Maximum Number of Cycles"]
letters = ["a)", "b)", "c)", "d)", "e)", "f)"]
for i in range(3):
    plt.subplot(2,3,i+1)
    sns.swarmplot(x="condition", y=metrics[i], data=phom,
                   palette=palette, order=["Awake", "Ketam", "Propo"])
    sns.boxplot(x="condition", y=metrics[i], data=phom,
                   palette=light_palette, order=["Awake", "Ketam", "Propo"])
    plt.title(titles[i])
    plt.ylabel("")
    plt.xlabel("Condition")
    plt.text(2.1, phom[metrics[i]].max(), letters[i], fontsize="x-large", fontweight="bold")
    
plt.subplot(2,3,4)
plt.scatter(phom_awake["max_pers"], phom_awake["ncycle"], color=palette[0], s=10)
plt.scatter(phom_ketam["max_pers"], phom_ketam["ncycle"], color=palette[1], s=10)
plt.scatter(phom_propo["max_pers"], phom_propo["ncycle"], color=palette[2], s=10)
plt.xlabel("Maximum Persistence")
plt.ylabel("Number of Cycles")
plt.text(0.625, 575, letters[3], fontsize="x-large", fontweight="bold")

plt.subplot(2,3,5)
plt.scatter(phom_awake["max_pers"], phom_awake["max_ncycles"], color=palette[0], s=10)
plt.scatter(phom_ketam["max_pers"], phom_ketam["max_ncycles"], color=palette[1], s=10)
plt.scatter(phom_propo["max_pers"], phom_propo["max_ncycles"], color=palette[2], s=10)
plt.xlabel("Maximum Persistence")
plt.ylabel("Maximum Number of Cycles")
plt.text(0.625, 130, letters[4], fontsize="x-large", fontweight="bold")

plt.subplot(2,3,6)
plt.scatter(phom_awake["ncycle"], phom_awake["max_ncycles"], color=palette[0], s=10)
plt.scatter(phom_ketam["ncycle"], phom_ketam["max_ncycles"], color=palette[1], s=10)
plt.scatter(phom_propo["ncycle"], phom_propo["max_ncycles"], color=palette[2], s=10)
plt.xlabel("Total Number of Cycles")
plt.ylabel("Maximum Number of Cycles")
plt.text(50, 130, letters[5], fontsize="x-large", fontweight="bold")

plt.tight_layout()
plt.savefig("phom_boxplots.pdf")
plt.show()

# %%
opn_awake = opn[opn["condition"] == "Awake"]
opn_ketam = opn[opn["condition"] == "Ketam"]
opn_propo = opn[opn["condition"] == "Propo"]

plt.figure(figsize=(12,8))
metrics = ["nodes", "edges", "determinism", "degeneracy", "modularity"]
titles = ["Number of Nodes", "Number of Edges", "Determinism",
          "Degeneracy", "Modularity"]
letters = ["a)", "b)", "c)", "d)", "e)"]
for i in range(5):
    plt.subplot(2,3,i+1)
    sns.violinplot(x="condition", y=metrics[i], data=opn,
                   palette=light_palette, order=["Awake", "Ketam", "Propo"])
    plt.title(titles[i])
    plt.ylabel("")
    plt.xlabel("Condition")
    
    if i == 2:
        plt.ylim([0.6,1])
        plt.text(2.1, 0.95, letters[i], fontsize="x-large", fontweight="bold")
    elif i == 3:
        plt.ylim([-0.01, 0.05])
        plt.text(2.1, 0.045, letters[i], fontsize="x-large", fontweight="bold")
    elif i == 4:
        plt.ylim([0.6,1.01])
        plt.text(2.1, 0.95, letters[i], fontsize="x-large", fontweight="bold")
    else:
        plt.text(2.1, opn[metrics[i]].max(), letters[i], fontsize="x-large", fontweight="bold")

plt.tight_layout()
plt.savefig("opn_violinplots.pdf")
plt.show()

# %%
plt.figure(figsize=(12,12))

#Nodes
plt.subplot(5, 5, 2)
plt.scatter(opn_awake["nodes"], opn_awake["edges"], color=light_palette[0], s=1)
plt.scatter(opn_ketam["nodes"], opn_ketam["edges"], color=light_palette[1], s=1)
plt.scatter(opn_propo["nodes"], opn_propo["edges"], color=light_palette[2], s=1)
plt.xlabel("Nodes")
plt.ylabel(("Edges"))

plt.subplot(5, 5, 3)
plt.scatter(opn_awake["nodes"], opn_awake["determinism"], color=light_palette[0], s=1)
plt.scatter(opn_ketam["nodes"], opn_ketam["determinism"], color=light_palette[1], s=1)
plt.scatter(opn_propo["nodes"], opn_propo["determinism"], color=light_palette[2], s=1)
plt.xlabel("Nodes")
plt.ylabel("Determinism")

plt.subplot(5, 5, 4)
plt.scatter(opn_awake["nodes"], opn_awake["degeneracy"], color=light_palette[0], s=1)
plt.scatter(opn_ketam["nodes"], opn_ketam["degeneracy"], color=light_palette[1], s=1)
plt.scatter(opn_propo["nodes"], opn_propo["degeneracy"], color=light_palette[2], s=1)
plt.xlabel("Nodes")
plt.ylabel("Degeneracy")

plt.subplot(5, 5, 5)
plt.scatter(opn_awake["nodes"], opn_awake["modularity"], color=light_palette[0], s=1)
plt.scatter(opn_ketam["nodes"], opn_ketam["modularity"], color=light_palette[1], s=1)
plt.scatter(opn_propo["nodes"], opn_propo["modularity"], color=light_palette[2], s=1)
plt.xlabel("Nodes")
plt.ylabel("Modulairty")

#Edges
plt.subplot(5, 5, 6)
plt.scatter(opn_awake["edges"], opn_awake["nodes"], color=light_palette[0], s=1)
plt.scatter(opn_ketam["edges"], opn_ketam["nodes"], color=light_palette[1], s=1)
plt.scatter(opn_propo["edges"], opn_propo["nodes"], color=light_palette[2], s=1)
plt.xlabel(("Edges"))
plt.ylabel("Nodes")

plt.subplot(5, 5, 8)
plt.scatter(opn_awake["edges"], opn_awake["determinism"], color=light_palette[0], s=1)
plt.scatter(opn_ketam["edges"], opn_ketam["determinism"], color=light_palette[1], s=1)
plt.scatter(opn_propo["edges"], opn_propo["determinism"], color=light_palette[2], s=1)
plt.xlabel(("Edges"))
plt.ylabel("Determinism")

plt.subplot(5, 5, 9)
plt.scatter(opn_awake["edges"], opn_awake["degeneracy"], color=light_palette[0], s=1)
plt.scatter(opn_ketam["edges"], opn_ketam["degeneracy"], color=light_palette[1], s=1)
plt.scatter(opn_propo["edges"], opn_propo["degeneracy"], color=light_palette[2], s=1)
plt.xlabel(("Edges"))
plt.ylabel("Degeneracy")

plt.subplot(5, 5, 10)
plt.scatter(opn_awake["edges"], opn_awake["modularity"], color=light_palette[0], s=1)
plt.scatter(opn_ketam["edges"], opn_ketam["modularity"], color=light_palette[1], s=1)
plt.scatter(opn_propo["edges"], opn_propo["modularity"], color=light_palette[2], s=1)
plt.xlabel(("Edges"))
plt.ylabel("Modulairty")

#Determinism 
plt.subplot(5, 5, 11)
plt.scatter(opn_awake["determinism"], opn_awake["nodes"], color=light_palette[0], s=1)
plt.scatter(opn_ketam["determinism"], opn_ketam["nodes"], color=light_palette[1], s=1)
plt.scatter(opn_propo["determinism"], opn_propo["nodes"], color=light_palette[2], s=1)
plt.xlabel("Determinism")
plt.ylabel("Nodes")

plt.subplot(5, 5, 12)
plt.scatter(opn_awake["determinism"], opn_awake["edges"], color=light_palette[0], s=1)
plt.scatter(opn_ketam["determinism"], opn_ketam["edges"], color=light_palette[1], s=1)
plt.scatter(opn_propo["determinism"], opn_propo["edges"], color=light_palette[2], s=1)
plt.xlabel("Determinism")
plt.ylabel(("Edges"))

plt.subplot(5, 5, 14)
plt.scatter(opn_awake["determinism"], opn_awake["degeneracy"], color=light_palette[0], s=1)
plt.scatter(opn_ketam["determinism"], opn_ketam["degeneracy"], color=light_palette[1], s=1)
plt.scatter(opn_propo["determinism"], opn_propo["degeneracy"], color=light_palette[2], s=1)
plt.xlabel("Determinism")
plt.ylabel("Degeneracy")

plt.subplot(5, 5, 15)
plt.scatter(opn_awake["determinism"], opn_awake["modularity"], color=light_palette[0], s=1)
plt.scatter(opn_ketam["determinism"], opn_ketam["modularity"], color=light_palette[1], s=1)
plt.scatter(opn_propo["determinism"], opn_propo["modularity"], color=light_palette[2], s=1)
plt.xlabel("Determinism")
plt.ylabel("Modulairty")

#Degeneracy
plt.subplot(5, 5, 16)
plt.scatter(opn_awake["degeneracy"], opn_awake["nodes"], color=light_palette[0], s=1)
plt.scatter(opn_ketam["degeneracy"], opn_ketam["nodes"], color=light_palette[1], s=1)
plt.scatter(opn_propo["degeneracy"], opn_propo["nodes"], color=light_palette[2], s=1)
plt.xlabel("Degeneracy")
plt.ylabel("Nodes")

plt.subplot(5, 5, 17)
plt.scatter(opn_awake["degeneracy"], opn_awake["edges"], color=light_palette[0], s=1)
plt.scatter(opn_ketam["degeneracy"], opn_ketam["edges"], color=light_palette[1], s=1)
plt.scatter(opn_propo["degeneracy"], opn_propo["edges"], color=light_palette[2], s=1)
plt.xlabel("Degeneracy")
plt.ylabel(("Edges"))

plt.subplot(5, 5, 18)
plt.scatter(opn_awake["degeneracy"], opn_awake["determinism"], color=light_palette[0], s=1)
plt.scatter(opn_ketam["degeneracy"], opn_ketam["determinism"], color=light_palette[1], s=1)
plt.scatter(opn_propo["degeneracy"], opn_propo["determinism"], color=light_palette[2], s=1)
plt.xlabel("Degeneracy")
plt.ylabel("Determinism")

plt.subplot(5, 5, 20)
plt.scatter(opn_awake["degeneracy"], opn_awake["modularity"], color=light_palette[0], s=1)
plt.scatter(opn_ketam["degeneracy"], opn_ketam["modularity"], color=light_palette[1], s=1)
plt.scatter(opn_propo["degeneracy"], opn_propo["modularity"], color=light_palette[2], s=1)
plt.xlabel("Degeneracy")
plt.ylabel("Modulairty")

#Modularity
plt.subplot(5, 5, 21)
plt.scatter(opn_awake["modularity"], opn_awake["nodes"], color=light_palette[0], s=1)
plt.scatter(opn_ketam["modularity"], opn_ketam["nodes"], color=light_palette[1], s=1)
plt.scatter(opn_propo["modularity"], opn_propo["nodes"], color=light_palette[2], s=1)
plt.xlabel("Modularity")
plt.ylabel("Nodes")

plt.subplot(5, 5, 22)
plt.scatter(opn_awake["modularity"], opn_awake["edges"], color=light_palette[0], s=1)
plt.scatter(opn_ketam["modularity"], opn_ketam["edges"], color=light_palette[1], s=1)
plt.scatter(opn_propo["modularity"], opn_propo["edges"], color=light_palette[2], s=1)
plt.xlabel("Modularity")
plt.ylabel(("Edges"))

plt.subplot(5, 5, 23)
plt.scatter(opn_awake["modularity"], opn_awake["determinism"], color=light_palette[0], s=1)
plt.scatter(opn_ketam["modularity"], opn_ketam["determinism"], color=light_palette[1], s=1)
plt.scatter(opn_propo["modularity"], opn_propo["determinism"], color=light_palette[2], s=1)
plt.xlabel("Modularity")
plt.ylabel("Determinism")

plt.subplot(5, 5, 24)
plt.scatter(opn_awake["modularity"], opn_awake["degeneracy"], color=light_palette[0], s=1)
plt.scatter(opn_ketam["modularity"], opn_ketam["degeneracy"], color=light_palette[1], s=1)
plt.scatter(opn_propo["modularity"], opn_propo["degeneracy"], color=light_palette[2], s=1)
plt.xlabel("Modularity")
plt.ylabel("Degeneracy")

plt.tight_layout()
plt.savefig("pairwise_comps.pdf")
plt.show()


# %%
plt.figure(figsize=(10,8))

plt.subplot(2,3,1)
sns.violinplot(x="condition", y="perm_ent", data=opn,
               palette=light_palette, order=["Awake", "Ketam", "Propo"])
plt.ylim([2,5.25])
plt.xlabel("Condition")
plt.title("Permutation Entropy")
plt.ylabel("")
plt.text(2, 4.9, "a)", fontsize="x-large", fontweight="bold")

plt.subplot(2,3,2)
sns.violinplot(x="condition", y="lyapunov", data=opn,
               palette=light_palette, order=["Awake", "Ketam", "Propo"])
plt.xlabel("Condition")
plt.title("Lyapunov Exponent")
plt.ylabel("")
plt.text(2, 0.23, "b)", fontsize="x-large", fontweight="bold")

plt.subplot(2,3,3)
plt.scatter(opn_awake["perm_ent"], opn_awake["lyapunov"], color=light_palette[0], s=1)
plt.scatter(opn_ketam["perm_ent"], opn_ketam["lyapunov"], color=light_palette[1], s=1)
plt.scatter(opn_propo["perm_ent"], opn_propo["lyapunov"], color=light_palette[2], s=1)
plt.xlabel("Permutation Entropy")
plt.ylabel("Lyapunov Exponent")
plt.text(0.5, 0.22, "c)", fontsize="x-large", fontweight="bold")

plt.subplot(2,3,4)
plt.scatter(opn_awake["perm_ent"], opn_awake["nodes"], color=light_palette[0], s=1)
plt.scatter(opn_ketam["perm_ent"], opn_ketam["nodes"], color=light_palette[1], s=1)
plt.scatter(opn_propo["perm_ent"], opn_propo["nodes"], color=light_palette[2], s=1)
plt.xlabel("Permutation Entropy")
plt.ylabel("Number of Nodes")
plt.text(0.5, 110, "d)", fontsize="x-large", fontweight="bold")


plt.subplot(2,3,5)
plt.scatter(opn_awake["lyapunov"], opn_awake["determinism"], color=light_palette[0], s=1)
plt.scatter(opn_ketam["lyapunov"], opn_ketam["determinism"], color=light_palette[1], s=1)
plt.scatter(opn_propo["lyapunov"], opn_propo["determinism"], color=light_palette[2], s=1)
plt.xlabel("Lyapunov Exponent")
plt.ylabel("Determinism")
plt.text(0.22, 0.925, "e)", fontsize="x-large", fontweight="bold")

plt.subplot(2,3,6)
plt.scatter(opn_awake["lyapunov"], opn_awake["modularity"], color=light_palette[0], s=1)
plt.scatter(opn_ketam["lyapunov"], opn_ketam["modularity"], color=light_palette[1], s=1)
plt.scatter(opn_propo["lyapunov"], opn_propo["modularity"], color=light_palette[2], s=1)
plt.xlabel("Lyapunov Exponent")
plt.ylabel("Modularity")
plt.text(0.225, 0.8, "f)", fontsize="x-large", fontweight="bold")


plt.tight_layout()
plt.savefig("timseries_analysis.pdf")
# %%
```
