## Supplementary material for "Topological Analysis of Differential Effects of Ketamine and Propofol Anesthesia on Brain Dynamics": scripts and csv files: filter.py

```
#!/usr/bin/env python3
# -*- coding: utf-8 -*-
"""
Created on Mon Jun  3 16:44:12 2019

@author: thosvarley
"""
import numpy as np 
from scipy.signal import decimate
import networkx as nx 
import os 
from pathlib import Path
import mne

cwd = "/script/dir/"
ptd = str(Path(cwd).parent)
in_dir = ptd + "/processed/raw"
out_dir = ptd + "/processed/filtered"

for i in os.listdir(in_dir):
    name = i[:-4]
    timeseries = np.load(in_dir + "/" + i)["arr_0"].astype("float32")
    
    ch_names = [str(x) for x in range(len(timeseries))]
    ch_types = ["ecg" for x in range(len(timeseries))]
    sfreq = 1000
    info = mne.create_info(ch_names=ch_names, sfreq=sfreq, ch_types = ch_types)
    picks = mne.pick_types(info, meg=False, ecg = True)
    
    timeseries_raw = mne.io.RawArray(timeseries, info)
    
    timeseries_raw.filter(l_freq = 0.1, h_freq = None, phase="zero-double", picks=picks) #High pass filter
    timeseries_raw.filter(l_freq = None, h_freq = 200, phase="zero-double", picks=picks)
    timeseries_raw.notch_filter(np.arange(50, 250, 50), picks=picks, phase="zero-double")
    
    timeseries = timeseries_raw._data
    
    #timeseries = decimate(timeseries, 2)
    
    mean = np.mean(timeseries)
    std = np.std(timeseries)
    
    timeseries = (timeseries - mean)/std
    
    timeseries = timeseries[:,50:-50]
    np.save(out_dir + "/{0}_filtered".format(name), timeseries)
    
    
```
