## Supplementary material for "Topological Analysis of Differential Effects of Ketamine and Propofol Anesthesia on Brain Dynamics": scripts and csv files: make_cuts.py

```
#!/usr/bin/env python3
# -*- coding: utf-8 -*-
"""
Created on Fri May 31 15:22:49 2019

@author: thosvarley
"""
import numpy as np 

from pathlib import Path
home = "/home/thosvarley/Data/neurotycho/anes_sleep/"

cwd = '/home/thosvarley/Desktop/cuts_stationary/scripts'
ptd = str(Path(cwd).parent)
out_dir = ptd + "/processed/raw"

PF730_Awake = "20120730PF_Anesthesia+and+Sleep_Chibi_Toru+Yanagawa_mat_ECoG128/Session1/raw_data/"
PF730_Propo = "20120730PF_Anesthesia+and+Sleep_Chibi_Toru+Yanagawa_mat_ECoG128/Session2/raw_data/"

PF731_Awake = "20120731PF_Anesthesia+and+Sleep_George_Toru+Yanagawa_mat_ECoG128/Session1/raw_data/"
PF731_Propo = "20120731PF_Anesthesia+and+Sleep_George_Toru+Yanagawa_mat_ECoG128/Session2/raw_data/"

PF802_Awake = "20120802PF_Anesthesia+and+Sleep_Chibi_Toru+Yanagawa_mat_ECoG128/Session1/raw_data/"
PF802_Propo = "20120802PF_Anesthesia+and+Sleep_Chibi_Toru+Yanagawa_mat_ECoG128/Session2/raw_data/"

PF803_Awake = "20120803PF_Anesthesia+and+Sleep_George_Toru+Yanagawa_mat_ECoG128/Session1/raw_data/"
PF803_Propo = "20120803PF_Anesthesia+and+Sleep_George_Toru+Yanagawa_mat_ECoG128/Session2/raw_data/"

KT719_Awake = "20120719KT_Anesthesia+and+Sleep_Chibi_Toru+Yanagawa_mat_ECoG128/Session1/raw_data/"
KT719_Ketam = "20120719KT_Anesthesia+and+Sleep_Chibi_Toru+Yanagawa_mat_ECoG128/Session2/"

KT724_Awake = "20120724KT_Anesthesia+and+Sleep_George_Toru+Yanagawa_mat_ECoG128/Session1/"
KT724_Ketam = "20120724KT_Anesthesia+and+Sleep_George_Toru+Yanagawa_mat_ECoG128/Session2/"

KT810_Awake = "20120810KT_Anesthesia+and+Sleep_George_Toru+Yanagawa_mat_ECoG128/Session1/"
KT810_Ketam = "20120810KT_Anesthesia+and+Sleep_George_Toru+Yanagawa_mat_ECoG128/Session2/"

KT813_Awake = "20120813KT_Anesthesia+and+Sleep_Chibi_Toru+Yanagawa_mat_ECoG128/Session1/"
KT813_Ketam = "20120813KT_Anesthesia+and+Sleep_Chibi_Toru+Yanagawa_mat_ECoG128/Session2/"

dirs = [PF730_Awake, PF730_Propo, 
        PF731_Awake, PF731_Propo, 
        PF802_Awake, PF802_Propo,
        PF803_Awake, PF803_Propo,
        KT719_Awake, KT719_Ketam,
        KT724_Awake, KT724_Ketam,
        KT810_Awake, KT810_Ketam,
        KT813_Awake, KT813_Ketam]

PF730_Awake_cuts = [("0049","0054"),("0100","0105"),("0116","0121")]
PF730_Propo_cuts = [("0556","0561"),("0586","0591"),("0600","0605"),("0615","0620"),("0630","0635"),("0641","0646")]
 
PF731_Awake_cuts = [("0001","0006"),("0012","0017"),("0070","0075")]
PF731_Propo_cuts = [('0787', '0792'), ('0805', '0810'), ('0826', '0831'), ('0844', '0849'), ('0878', '0883'), ('0893', '0898')]

PF802_Awake_cuts = [('0063','0068'),('0085','0090'),('0107','0112')]
PF802_Propo_cuts = [('0735', '0740'), ('0746', '0751'), ('0774', '0779'), ('0799', '0804'), ('0820', '0825'), ('0831', '0836')]

PF803_Awake_cuts = [('0015','0020'),('0029','0034'),('0077','0082')]
PF803_Propo_cuts = [('0618', '0623'),('0630', '0635'),('0658', '0663'),('0684', '0689'),('0696', '0701'),('0710', '0715')]

KT719_Awake_cuts = [('0380','0385'),('0391','0396'),('0418','0423')]
KT719_Ketam_cuts = [('0932', '0937'),('0958', '0963'),('0986', '0991'),('999', '1004'),('1020', '1025'),('1031', '1036')]

KT724_Awake_cuts = [('0085','0090'),('0139','0144'),('0260','0265')]
KT724_Ketam_cuts = [('1355', '1360'),('1370', '1375'),('1390', '1395'),('1415', '1420'),('1430', '1435'),('1443', '1448')]

KT810_Awake_cuts = [('0001','0006'),('0019','0024'),('0041','0046')]
KT810_Ketam_cuts = [('0366', '0371'),('0380', '0385'),('0395', '0400'),('0415', '0420'),('0430', '0435'),('0450', '0455')]

KT813_Awake_cuts = [('0028','0033'),('0039','0044'),('0054','0059')] #5 sec
KT813_Ketam_cuts = [('0677', '0682'),('0699', '0704'),('0732', '0737'),('0775', '0780'),('0789', '0794'),('0811', '0816')]

cuts = [PF730_Awake_cuts, PF730_Propo_cuts, 
        PF731_Awake_cuts, PF731_Propo_cuts, 
        PF802_Awake_cuts, PF802_Propo_cuts,
        PF803_Awake_cuts, PF803_Propo_cuts,
        KT719_Awake_cuts, KT719_Ketam_cuts,
        KT724_Awake_cuts, KT724_Ketam_cuts,
        KT810_Awake_cuts, KT810_Ketam_cuts,
        KT813_Awake_cuts, KT813_Ketam_cuts]

cuts_str = ["PF730_Awake", "PF730_Propo", 
            "PF731_Awake", "PF731_Propo", 
            "PF802_Awake", "PF802_Propo",
            "PF803_Awake", "PF803_Propo",
            "KT719_Awake", "KT719_Ketam",
            "KT724_Awake", "KT724_Ketam",
            "KT810_Awake", "KT810_Ketam",
            "KT813_Awake", "KT813_Ketam"]

chibi_bad_channels = (22,23,24,33)
george_bad_channels = (34,73)

for i in range(len(dirs)):
    ecog = np.load(home + dirs[i] + "dataset_raw.npy", allow_pickle=True)[0]
    cut = cuts[i]
    
    for j in cuts[i]:
        start = int(j[0])
        end = int(j[1])
        ecog_cut = ecog[:,(start*1000):(end*1000)]
        
        if "Chibi" in dirs[i]:
            ecog_cut = np.delete(ecog_cut, chibi_bad_channels, axis=0)
        elif "George" in dirs[i]:
            ecog_cut = np.delete(ecog_cut, george_bad_channels, axis=0)
        
        np.save(out_dir + "/" + cuts_str[i] + "_" + j[0] + "_" + j[1] + ".npy", ecog_cut)
        
    
    
```
