## Supplementary material for "Topological Analysis of Differential Effects of Ketamine and Propofol Anesthesia on Brain Dynamics": scripts and csv files: morphospace.py

```
#!/usr/bin/env python3
# -*- coding: utf-8 -*-
"""
Created on Thu Aug 13 15:46:37 2020

@author: thosvarley
"""
import pandas as pd 
import numpy as np 
import matplotlib.pyplot as plt
import seaborn as sns 

from sklearn.preprocessing import StandardScaler
from sklearn.decomposition import PCA
from umap import UMAP
from mpl_toolkits.mplot3d import Axes3D
from copy import deepcopy

sns.set(style="darkgrid")
palette = sns.color_palette(n_colors=3, palette="husl")

in_dir = "/home/thosvarley/Desktop/cuts_stationary/results/"
out_dir = '/home/thosvarley/Desktop/cuts_stationary/'

results = pd.read_csv(in_dir + "OPN_results.csv", index_col = "Unnamed: 0")
opns = deepcopy(results)
awake = results[results["condition"] == "Awake"]
ketam = results[results["condition"] == "Ketam"]
propo = results[results["condition"] == "Propo"]

columns = ["nodes", "edges", "determinism", 
           "degeneracy", "modularity"]

measures = results[columns]
conditions = np.array([x[6:11] for x in measures.index])
awake_coords = np.where(conditions == "Awake")[0]
ketam_coords = np.where(conditions == "Ketam")[0]
propo_coords = np.where(conditions == "Propo")[0]

ss = StandardScaler()
measures_ss = ss.fit_transform(measures.values)

pca = PCA(n_components=3)
measures_pca = pca.fit_transform(measures_ss)

umap = UMAP()
measures_umap = umap.fit_transform(measures_ss)

plt.figure(figsize=(8,6))

plt.subplot(1,2,1)
plt.scatter(measures_ss[awake_coords,5], measures_ss[awake_coords,-1],
            c=palette[0], s=5, label="Awake")
plt.scatter(measures_ss[ketam_coords,5], measures_ss[ketam_coords,-1],
            c=palette[1], s=5, label="Ketamine")
plt.scatter(measures_ss[propo_coords,5], measures_ss[propo_coords,-1],
            c=palette[2], s=5, label="Propofol")
plt.xlabel("Modularity")
plt.ylabel("Permutation Entropy")
plt.title("Modularity/Entropy")
plt.legend()

plt.subplot(1,2,2)

plt.scatter(measures_umap[awake_coords,0], measures_umap[awake_coords,1],
            c=palette[0], s=5, label="Awake")
plt.scatter(measures_umap[ketam_coords,0], measures_umap[ketam_coords,1],
            c=palette[1], s=5, label="Ketamine")
plt.scatter(measures_umap[propo_coords,0], measures_umap[propo_coords,1],
            c=palette[2], s=5, label="Propofol")
plt.xlabel("Axis 1")
plt.ylabel("Axis 2")
plt.title("Ordinal Partition Network Morphospace Embedding")
plt.legend()

plt.tight_layout()
plt.savefig("opn_umap.pdf", dpi=250)


#DENSITY, NOT EDGE COUNT 
opns["density"] = opns["edges"] / (opns["nodes"]**2)

plt.figure(figsize=(8,4))
plt.subplot(1,2,1)
sns.violinplot(x="condition", y="determinism", data=opns, palette=palette)
plt.xticks([0,1,2], ["Awake", "Ketamine", "Propofol"])
plt.xlabel("Condition")
plt.ylabel("Bit")
plt.ylim([0.6,1])
plt.title("Determinism***")

plt.subplot(1,2,2)
sns.violinplot(x="condition", y="degeneracy", data=opns, palette=palette)
plt.xticks([0,1,2], ["Awake", "Ketamine", "Propofol"])
plt.xlabel("Condition")
plt.ylabel("Bits")
plt.title("Degeneracy***")
plt.ylim([-0.01,0.05])

plt.tight_layout()
plt.savefig(out_dir + "det_deg.png", dpi=250)
plt.show()

plt.figure()
sns.violinplot(x="condition", y="modularity", data=opns, palette=palette)
plt.ylim([0.6,1])
plt.xticks([0,1,2], ["Awake", "Ketamine", "Propofol"])
plt.xlabel("Condition")
plt.ylabel("Q")
plt.title("Modularity***")
plt.savefig(out_dir + "modularity.png", dpi=250)
```
