## Supplementary material for "Topological Analysis of Differential Effects of Ketamine and Propofol Anesthesia on Brain Dynamics": S.I. Figures

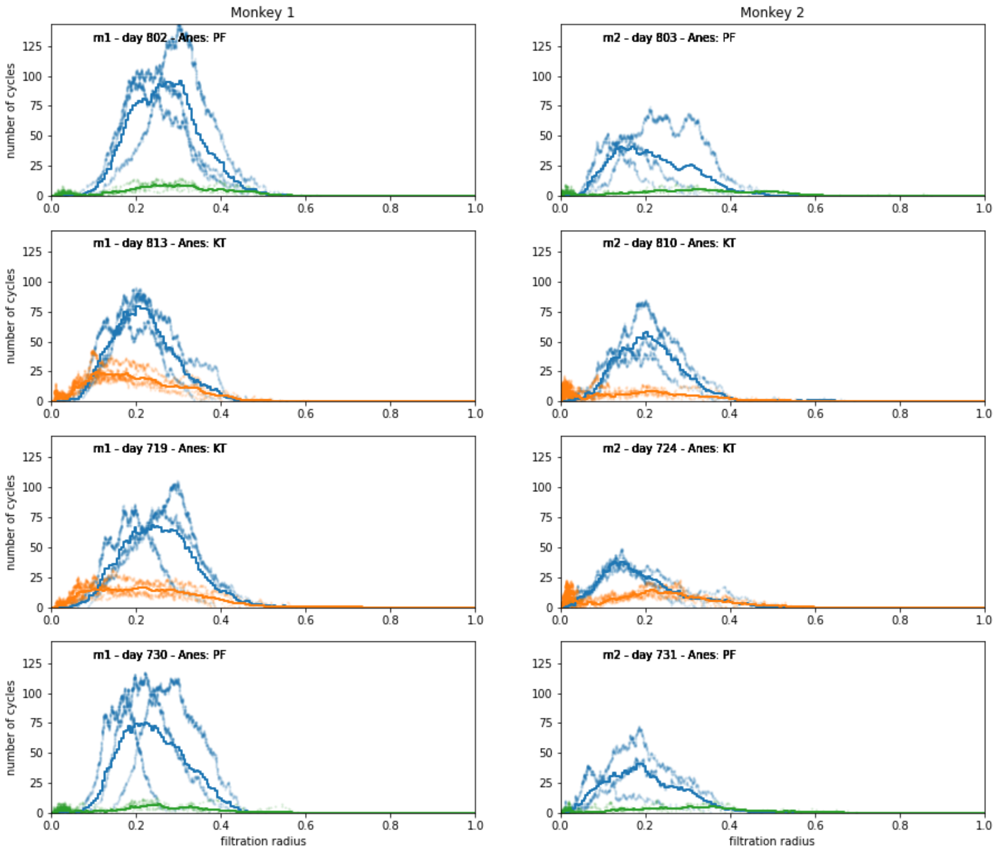


Figure 1: Average Betti curves divided by monkeys(columns) and by days (rows). The curve represent the average number of cycles present at the corresponding value of the filtration. Note that different states are largely consistent between monkeys, but very different from each other. BLUE: Awake, ORANGE: Ketamine, GREEN: Propofol.

### References

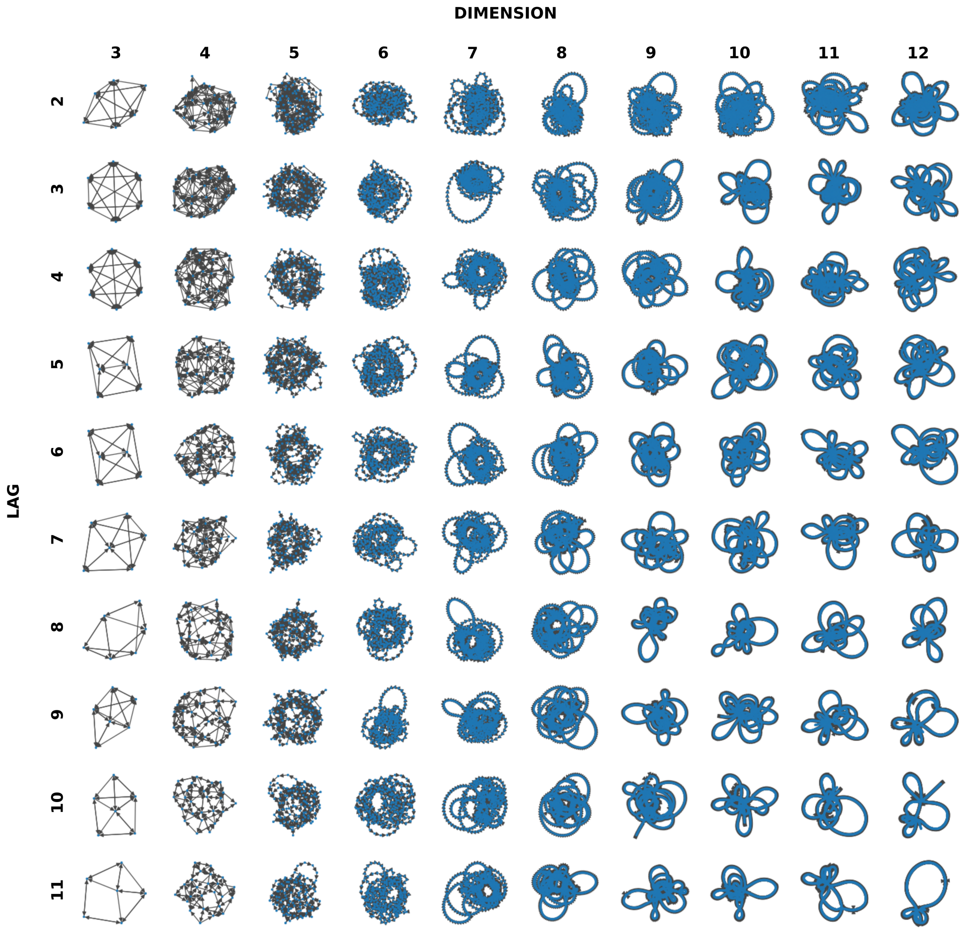


Figure 2: Example that shows the effect of the lag and dimension parameters on the creation of an OPN network. Each row corresponds to a lag, and each column to an embedding dimension: note that as dimension and lag increase, eventually the associated OPN becomes increasingly dominated by paths of unique nodes.

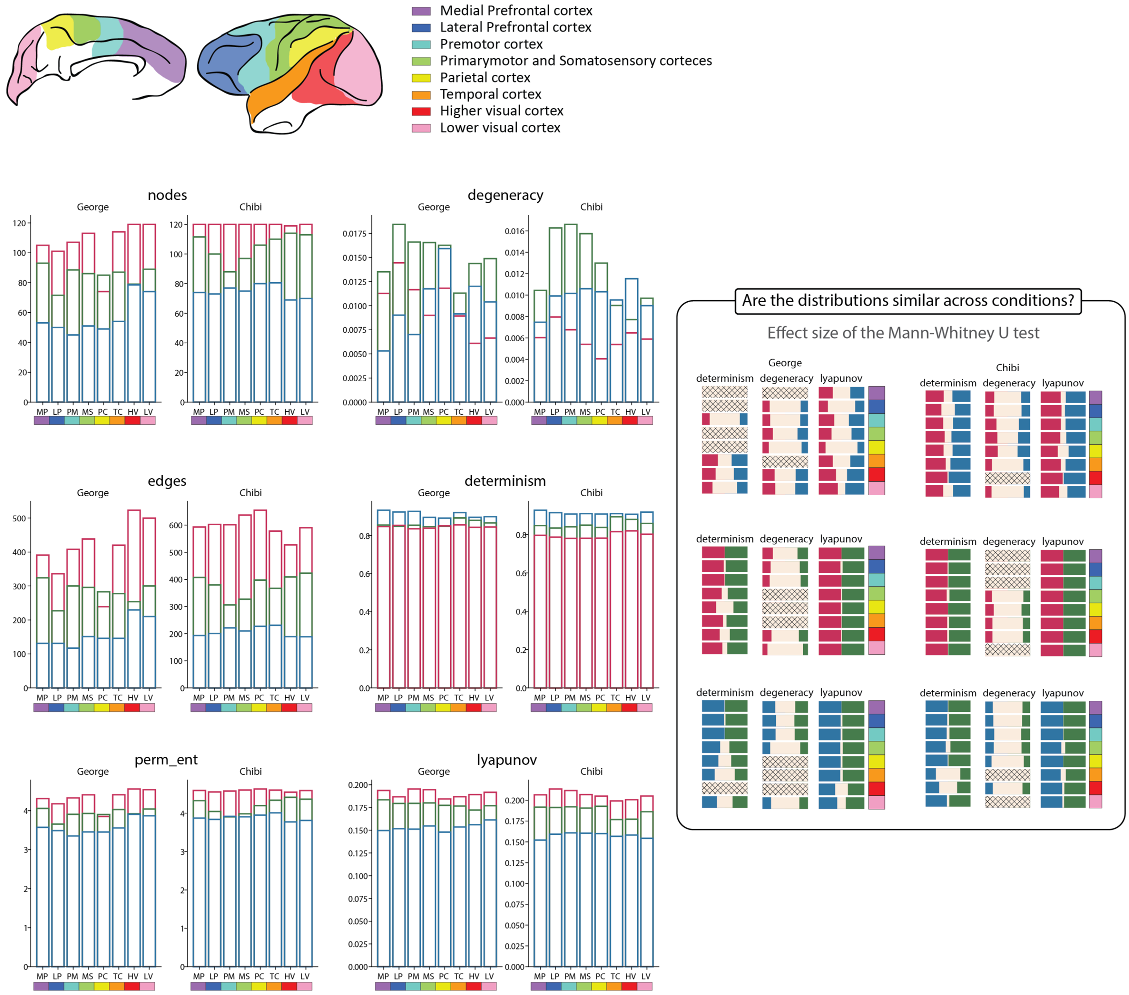


Figure 3: (left) Bar plots representing the median of the samples for 6 statistics relative to the OPNs divided by anatomical region. (right) The effect size measure goes from 0 to .5 . For 0, the two distributions are completely separated, for .5 the two distributions completely overlap. In this image white band represents the effect size, the smaller the band the less the two distributions overlap (GREEN: ketamine, BLUE: propofol, PINK: awake). The patterned bars represent the comparisons for which we accept the null hypothesis of the test and the two distributions are significantly similar.

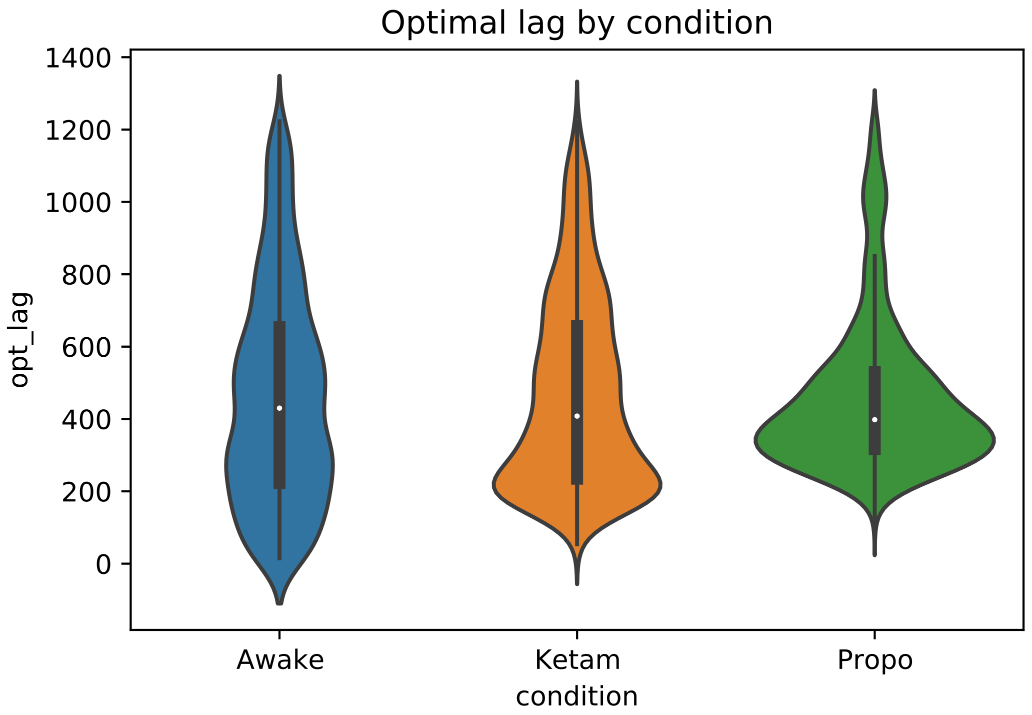


Figure 4: The distribution of optimal lags between the three conditions. The differences between conditions were extremely small: the ketamine condition technically had the longest "memory", with an optimal lag of $469.68 \pm 267.47$, followed by the awake condition ($465.02 \pm 301.98$ and then the propofol condition with $459.76 \pm 210.63$.
